# Directed evolution for cell separation in natural isolates of budding yeast reveals selection to deactivate *AMN1* and the Rim101 pathway in haploids and selection in favor of Hawthorne’s deletion in diploids

**DOI:** 10.1101/2024.05.03.592462

**Authors:** Benjamin Galeota-Sprung, Erik Pritchard, Crystal Huang, Amy Fernandez, Paul Sniegowski

## Abstract

Natural isolates of the yeast *Saccharomyces cerevisiae* were evolved under a transfer protocol that selected for cell separation and against clumpy growth. Whole-genome sequencing of haploid populations revealed strong selection to deactivate *AMN1*, a known regulator of post-mitotic cell separation, as well as multiple instances of loss-of-function mutations on the Rim101 pathway, pointing to a previously unknown role of the Rim101 pathway in regulating cell separation. In diploid populations, we observed repeated large partial deletions of chromosome III caused by fusions of the mating type loci *MAT* and HMR (Hawthorne’s deletion) or *MAT* and *HML* (Strathern’s circle). We measured the spontaneous rate of Hawthorne’s deletion and found that it is within an order of magnitude of previously measured rates of whole-chromosome aneuploidy. A diploid population in which neither large deletion was detected instead fixed a heterozygous nonsynonymous mutation to the calcium channel *CCH1*, also pointing to a novel role for this gene in relation to cell separation.

## Introduction

Yeasts are fungi that have re-evolved unicellularity as a derived trait from a multicellular fungal ancestor. Although they are unicellular, yeasts such as *S. cerevisiae* are capable of phenotypic states in which individual cells are in sustained contact with one another. These various cell aggregation phenotypes include mating, flocculation, pseudohyphal growth, invasive growth, biofilms, and clumpy growth (Soares 2011). The latter phenomenon, clumpy growth (also called chain formation (Stewart 2009)), occurs when growing cells do not fully separate after the mitotic process of budding, leading to clusters of related cells. The fundamental importance of the regulation of post-mitotic separation to both pathogenicity (*i.e*. biofilms), and to organismal complexity and multicellularity, has led to sustained interest in experimental selection for decreased post-mitotic separation (Ratcliff et al. 2015; Hope et al. 2017; Tong et al. 2025).

Typical laboratory strains of *S. cerevisiae* are not clumpy. Much of the variation in clumpiness across strains is explained by different alleles of the cell separation regulator *AMN1* (Li et al. 2013), which represses the cell separation program in haploids but not diploids (Fang et al. 2018). Laboratory strains such as S288C have a nonfunctional allele of *AMN1*. Since cultures of well-separated cells are easier to work with in many respects, this phenotype is advantageous to the experimenter and may have been semi-inadvertently selected for in the domestication history of common laboratory strains. In contrast to laboratory strains, natural isolates (“wild strains”) of *S. cerevisiae* are often clumpy growers, especially as haploids (Figure 1B). Wild strains tend to be more robust than lab strains in many respects, including faster growth rate, higher sporulation efficiency, and lower rate of petite production (Dimitrov et al., 2009). These properties of wild strains are particularly interesting given the many industrial uses of *S. cerevisiae*. Wild strains are also interesting in light of the status of *S. cerevisiae* as an important model organism coupled with the well-known, but not necessarily well-characterized, importance of strain-specific background effects on genotype-phenotype mappings (e.g. (Galardini et al. 2019)). While some properties of wild strains render them more experimentally tractable than common laboratory strains, their clumpy growth presents technical difficulties for otherwise routine assays such as dilution plating to estimate population density and flow cytometry of single cells.

**Figure 1.**
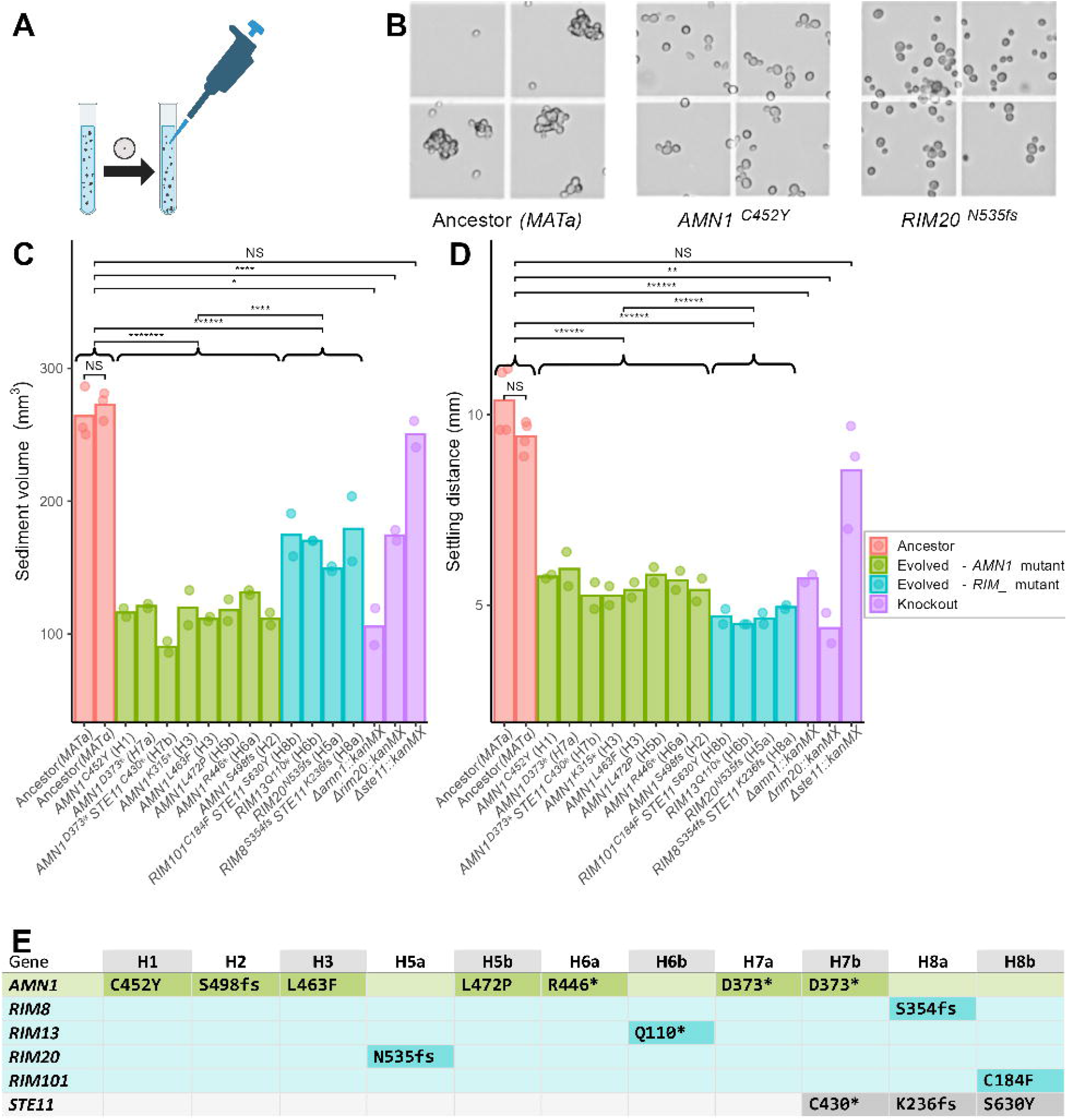
(A) Large clusters were allowed to settle out prior to transfer. (B) The clumpy phenotype of the haploid ancestor compared to the separated phenotype of representative evolved clones. (C) Evolved and knockout haploid strains have less sediment at the bottom of a culture after 2 h of settling. (D) Evolved and knockout haploid strains have less downwards progress of the cloudy area of culture after 2h settling. Curly braces indicate group comparisons. D and E share the same legend. (E) Across clones isolated from 11 haploid populations, parallel changes at the gene (*AMN1, STE11*) or pathway (Rim101) level were observed.

We were motivated to fill a gap in the literature by performing experimental selection for increased cell separation. Because they are clumpier to begin with, and because we were interested in creating less clumpy variants of wild strains, we carried out this experiment using wild strains of *S. cerevisiae*. In light of the role of *AMN1* as a primary regulator of post-mitotic separation in haploids, we founded populations with both haploid and diploid ancestors. We evolved replicate populations for 57 days (>380 generations) under a daily transfer protocol that selected against large clumps and for separated cells. We report here the genotypic and phenotypic evolution observed in this experiment.

## Methods

### Medium and growth conditions

The evolution experiment was carried out in glass 50 mL Erlenmeyer-type flasks containing 10 mL synthetic defined (SD) medium (6.7 g/L yeast nitrogen base, 2% glucose) supplemented with ampicillin and tetracycline, at 30 C and shaken at 200 rpm. SD was chosen as the medium for the evolution experiment because cultures of the wild diploid strains used are already very well-separated in YPD after the exponential growth phase. For haploid clones isolated from the evolution experiment, some assays were carried out in YPD (2% peptone, 1% yeast extract, 2% glucose).

### Strains

The diploid strain YPS602 is a monosporic isolate of YPS133, which was isolated from Tyler Arboretum, Pennsylvania, as described previously (Sniegowski et al. 2002). The closely related diploid strain YPS606 is a monosporic isolate of YPS142, also isolated from Tyler Arboretum. Both strains are related to the more widely known YPS128, and are members of the North American Oak clade (Peter et al. 2018). YPS2070 is a haploid *MATα* heterothallic *ho::kanMX* derivative of YPS602 and YPS2055 is a haploid *MAT*a heterothallic *ho::kanMX* derivative of YPS606.

### Evolution experiment

Four replicate populations of each of the 4 strains described above were founded from single colonies and transferred daily according to the following protocol (illustrated schematically in Figure 1A). A 2 mL aliquot of well-mixed culture was removed to a small glass tube and allowed to settle undisturbed for 10 min (Day 1 – Day 8), 20 min (Day 9 – Day 14), or 30 min (Day 15 – Day 57). After the waiting period, 100 µL of culture was carefully removed from near the top of the aliquot and transferred to 9.9 mL fresh medium in a new flask.

Transfers were performed every ∼24h for 57 days (implying at least 57 × log_2_100 = 380 generations). At regular intervals, 3 mL of culture was frozen in 15% glycerol and stored at −80 C. Clonal isolates were obtained from frozen population samples by streaking to single colonies. For a control (no selection against clumping) evolution experiment, conditions were identical except that 50 µl of well-mixed culture was added to 9.95 mL fresh medium at each transfer, for 66 days.

### Populations analyzed

The 16 populations founded from the haploids YPS2070 and YPS2055 and the diploids YPS602 and YPS606 were designated H1-H4, H5-H8, D1-D4, and D5-D8, respectively. After completion of the evolution experiment, it became apparent that at some point between days 29 and 35, the diploid populations D5-D8 were inadvertently discontinued while haploid populations H5-H8 were duplicated. We subsequently adopted an *a/b* naming scheme to distinguish these duplicated populations. Population H4 was lost before the end of the experiment due to a separate technical error. Therefore after 57 days of evolution there were 11 haploid populations (H1, H2, H3, H5a, H5b, H6a, H6b, H7a, H7b, H8a, H8b) and 4 diploid populations (D1, D2, D3, D4) available for analysis, and additionally the diploid populations D5-D8 completed 29 days of evolution.

### Settling and sedimentation assays

When a well-mixed culture of yeast is placed in a clear vessel and allowed to settle, two phenomena are apparent: (1) sediment collects at the bottom of the vessel and (2) the cloudy area of culture suspension moves visibly downwards, with clear medium above. We measured both manifestations of settling (for illustrative photographs see Figure S2 in File S1). Haploid assays were performed in YPD while diploid assays were performed in SD. For all assays, clones were started from a frozen chunk and grown overnight in YPD, and then grown for two 24h cycles in SD or one 24h cycle in YPD, diluting 1/100 at each transfer, prior to the assay. In the sedimentation assays, 5 mL of well-mixed culture was placed into a 15 mL conical tube suspended vertically; after two hours, the height of the sediment at the bottom was measured to the nearest tenth of a millimeter using a digital caliper tool. This height was subsequently converted to a volume. For haploids, the settling distance was measured concurrently with the sediment measurement, from the same 5 mL aliquot, estimated as the distance from the meniscus to the midpoint between clear media and cloudiest culture. For diploids, the settling distance was measured by a slightly different procedure: 900 μl of culture was briefly spun down (90 s at 10,000 rcf) and resuspended in 900 μl sterile water. Then 700 μl of this resuspension was pipetted into a narrow-profile cuvette and allowed to settle. We used cuvettes because we found that the flat surface aided in accurate measurement, and the resuspension step was necessary because cultures in SD do not settle evenly. After 2 hours, the downward progression of the visible demarcation from clear to cloudy was measured to the nearest tenth of a millimeter using a digital caliper tool.

### DNA extraction and whole-genome sequencing

We used the genome assembly of YPS128 as provided by Yue *et al*. (2017) as the reference for all genomic analyses described here. For clones, a culture was started from a frozen chunk, while for frozen populations, the entire frozen sample was thawed and 10 µl used to inoculate a new culture. After growth overnight in YPD, DNA was extracted from 1 mL of culture using kit D7005 from Zymo Research. Short-read whole-genome sequencing was performed on an Illumina NovaSeq X Plus sequencer in one or more multiplexed shared-flow-cell runs, producing 2×151bp paired-end reads. Reads were mapped to the reference genome using `bwa mem`(Li 2013), and duplicate reads removed by `PicardTools MarkDuplicates`. Mutations were called using `freebayes`(Garrison and Marth 2012) on the default settings, with all potential variants reviewed manually and examined in IGV (Robinson et al. 2011). Mutation calls from this pipeline were also spot-checked using `breseq`(Deatherage and Barrick 2014). A spreadsheet of all called mutations is available in the extended data. For the allele frequencies shown in Table 1, breseq was used in polymorphism mode, with a minimum allowed frequency of 5%. To locate chromosomal deletions, plots of read depth were created from the output of `samtools-depth`(Danecek et al. 2021). Additionally, long-read Nanopore sequencing was used to confirm the *MAT-HML* fusion in the clone isolated from population D2. To do so, we used `blastn`(Camacho et al. 2009) to map segments of genes bordering *HML* and *MAT* to all Nanopore reads >= 10 kb that mapped to chrIII, from which we detected single reads spanning both regions as evidence of fusion.

**Table 1.** Allele frequencies as revealed by whole population sequencing of two Day 57 haploid populations. Lineages with frequency <5% are not reported.

| Haploid population H3 |  | Haploid population H5a |  |
| --- | --- | --- | --- |
| Mutation | Freq | Mutation | Freq |
| *AMN1* K315* | 0.51 | *RIM20* N535fs | 0.42 |
| *AMN1* R162* | 0.13 | *AMN1* Q224* | 0.34 |
| *AMN1* E476* | 0.08 |  |  |
| *AMN1* L463F | 0.07 |  |  |

### Allele replacements

We replaced *AMN1* alleles in evolved clones with the ancestral *AMN1* sequence. To do so we inserted the *natMX* cassette just downstream of *AMN1* in the ancestral strain. We then amplified the entire *AMN1-natMX* cassette and transformed it into evolved strains. Successful transformation was confirmed by Sanger sequencing to ensure that the ancestral allele was introduced. Transformations were carried out by standard lithium acetate methods (Gietz and Schiestl 2007).

### Cluster size analysis

While microscopic inspection of evolved haploid strains showed a clear and obvious cell separation phenotype (Fig 1B), for diploids the phenotype was more subtle. To estimate the distribution of cluster sizes in diploid strains, we imaged and labelled growing cultures. A frozen chunk of the strain of interest was used to inoculate an overnight culture in YPD; the following day this culture was diluted 1/100 into fresh SD and allowed to grow for 24 h; then another 1/100 dilution and growth cycle in SD was performed; and finally on the 4^th^ day 200 µl of culture was transferred to 12 mL fresh SD and grown until A600=0.6, at which point photographs of the culture were taken at 100× magnification. Subsequently, the number of cells in each cluster was manually labelled (example shown in Figure S4 in file S1) and recorded for further analysis.

### Sporulation and tetrad dissection

Sporulation of diploid clones was induced by transferring an aliquot of saturated culture to 1% potassium acetate for incubation at 30 C for 3 days. 10 uL of sporulated culture was then added to 50 uL of a solution consisting of 0.5 mg/mL zymolyase (US Biological Z1005), 1 M sorbitol, and 0.1 M potassium phosphate buffer (pH 7.5) and incubated at 30 C for 10 minutes, after which 800 uL of sterile water was added to stop the digestion. A portion of the resulting solution was evenly spread onto an area of a YPD agar plate, and a light microscope equipped with a micromanipulator actuating a glass needle (Singer Instruments) was used to separate the tetrad.

### PCR verification of MAT-HMR and MAT-HML fusions

Primers 5’-TGGAAAGCGTAAACACGGAG-3’ and 5’-TCGGATTTGCGCTTGACAAT-3’ were used to verify the *MAT-HMR* deletion; these primers produce an amplicon of ∼3.5 kbp only in the case of a *MAT*-HMR fusion. Similarly, primers 5’-GTCCAGGGGCGGTTTATTTT-3’ and 5’-GGACTTGGAAGAAGCGTTGG-3’ were used to verify the MAT-HML fusion. Both primer sets were validated on known deletion strains as determined by whole-genome sequencing.

### Estimation of the spontaneous rate of Hawthorne’s deletion

We constructed the following heterothallic haploid strains in the YPS602 background. YPS4277 has the genotype *MATalpha ho::hphMX ura3::natMX his4::kanMX THR4*. YPS4263 has the genotype *MATa ho::hphMX ura3::natMX HIS4 thr4::URA3 amdSYM* with the *amdSYM* marker (Solis-Escalante et al. 2013) inserted on chrIII between *SED4* and *ATG15*. These strains were constructed by standard lithium acetate transformation methods. We then mated these strains creating the diploid YPS4283, which is sensitive to both 5-fluoroorotic acid (5-FOA) and fluoroacetamide (FAA). The markers responsible for the sensitivity were located on chrIII between *MAT* and HMR, so that a *MAT-HMR* deletion created a double 5-FOA/FAA resistant. Complete loss of the marked chrIII was selected against by the absence of histidine in growth media.

To estimate the rate of *MAT-HMR* fusion, we conducted a fluctuation assay (Gerrish 2008) with 4 replicate cultures. These cultures were inoculated from single colonies of YPS4283 into 6 mL SD + 20 mg/L uracil and grown overnight at 30 C with shaking, after which the density was estimated by hemocytometer and a target of 1000 cells transferred to 10 mL fresh SD + uracil. These sub-cultures were grown for 3 days at 30 C with shaking, and then 40 µL (1/250^th^ of the culture) was plated neat to SD + uracil + 1 g/L 5-FOA^R^ + 2.3 g/L FAA^R^ agar plates and the culture density estimated by hemocytometer counts. Plates were incubated for 2 days at 30 C and then examined. Very small colonies were excluded from subsequent analysis. The resulting colonies, being 5-FOA FAA His+, were considered likely to have the *MAT-HMR* fusion on the marked chrIII. (Nonsporulating colonies were likely *MAT-HML* fusions, but we did not measure this rate.) To confirm *MAT-HMR* fusions, we used a 2:2 live:dead segregation pattern of tetrads as a first check, and a PCR test as a second check. From each plate, 32-38 colonies were picked to both 1 mL YPD and to 1 mL SPO + uracil, in glass tubes. The YPD cultures were grown overnight and then stored at 4 C until needed. The SPO + uracil cultures were grown at 30 C for 3 days and then examined via microscope. From any culture in which sporulation was observed, at least 4 tetrads were dissected. If the dissection yielded a consistent pattern of 2:2 live:dead, DNA was extracted from the corresponding YPD culture and the presence of a *MAT-HMR* deletion was verified via PCR using the primers described above. Only one colony with a 2:2 live:dead segregation pattern did not give a positive PCR result (Table S2 in file S1). The total number of verified *MAT-HMR* deletions for each replicate was input into the package rSalvador (Zheng 2017), using the Lea-Coulson model with ε=1/250, to generate a point estimate and confidence interval for the spontaneous rate of *MAT-HMR* deletion.

### Protein alignment

The canonical amino acid sequence of human *CACNA1A* was downloaded from UniProt and aligned to *S. cerevisiae CCH1* using both Clustal Omega (Sievers et al. 2011) and TM-Coffee (Floden et al. 2016), a specialized aligner for transmembrane proteins. Both aligners agreed on the local alignment around the site of interest, yeast F994.

## Results

### All evolved haploid clones had a mutation to either AMN1 or the Rim101 pathway

We isolated one clone from each final haploid population, except in the case of population H3 we isolated two clones with different *AMN1* mutations. All 12 clones isolated from haploid Day 57 populations were markedly less clumpy when examined microscopically (examples shown in Figure 1B), and displayed changes of large and statistically significant magnitude in two assays measuring settling (Figs. 1C and 1D). Whole-genome sequencing revealed that all clones had either a mutation to *AMN1*, a known regulator of cell separation, or a gene in the Rim101 pathway: *RIM8, RIM13, RIM20*, or *RIM101*. There were no *AMN1* / Rim101 pathway double mutants.

### All observed AMN1 and Rim101 pathway mutations appeared to be loss-of-function

In total we observed 10 independent *AMN1* mutations (Figure 1E and Table 1). Of these, 7 are nonsense or frameshift while 3 are nonsynonymous amino acid substitutions. All 3 substitutions to *AMN1* occur between residues 452-472. Similarly, of the 4 mutations observed in the Rim101 pathway, three are frameshift or nonsense mutations and one is a substitution. Our supposition was that all *AMN1* and Rim101 pathway mutations, including the substitutions, were loss of function. Engineered *Δamn1* and *Δrim20* knockouts displayed similar phenotypes to the corresponding evolved clones (Figs. 1C and 1D). We also investigated two *AMN1* substitutions more closely: in the respective evolved clones, we replaced *AMN1*^*C452Y*^ and *AMN1*^*L472P*^ with the ancestral allele. These replacements completely restored the clumpy phenotype (Figure S1 in File S1), demonstrating that the substitutions alone were sufficient to cause the separated phenotype.

### The phenotype of AMN1 and Rim101 pathway mutants differs

While both *AMN1* and Rim101 pathway mutants are markedly less clumpy than the ancestor, settling assays reveal differences between the two genotypes. We allowed cultures to rest undisturbed for 2 h and measured two different aspects of the process of cells settling out of solution. Comparing group means, Rim101 pathway mutants accumulate more sediment at the bottom of the culture than do *AMN1* mutants (168.3 mm^3^ vs 114.9 mm^3^, p < 1e-04; Figure 1C). Conversely, in *AMN1* mutants, more downwards progress of the visibly cloudy area of culture (5.6 mm vs 4.7 mm, *p* < 1e-06; Figure 1D) is observed than in Rim101 pathway mutants.

### Haploid populations were diverse and experienced soft sweeps

Pairs of clones isolated from populations that were originally a single population tended to have different *AMN1* or Rim101 pathway mutations (Figure 1E noting *a* and *b* labels). This suggested that populations were diverse, with many contending lineages. To confirm this, we performed whole-population sequencing of haploid populations H3 and H5a. The results (Table 1) confirmed our hypothesis: we detected at least 6 independent *AMN1* or Rim101 pathway mutant lineages across the two populations. We suspect there are further low-frequency lineages that we were unable to resolve due to limitations of read depth.

### Parallel loss-of-function mutations to STE11 occurred in haploids

In the sequenced haploid clones we observed three independent mutations to *STE11*, each of which all appeared in conjunction with an *AMN1* or Rim101 pathway mutation. We confirmed that these clones are sterile (do not mate), as expected for loss-of-function mutations. Because populations H7a and H7b were once a single population and share the mutation *AMN1*^*D373\**^, we can infer with relative confidence that the *STE11*^*C430\**^ mutation arose in a background that already had fixed *AMN1*^*D373\**^. We surmise, then, that all observed *STE11* mutations were second mutations. Settling and sedimentation assays did not detect an effect of knocking out *STE11*; however, we did notice a subtle visual difference in settling appearance. It remains uncertain whether *ste11* mutations rose in frequency in response to the selection for cell separation, or whether they had a more general effect on fitness in the experiment.

### Large deletions involving the mating loci evolved multiple times in diploid populations

In contrast to haploid populations, in which all mutations affecting the cell separation phenotype that we detected were single-base-pair point mutations, in diploid populations we found that a large deletion on chromosome III evolved repeatedly (Figure 2A). This ∼90 kb deletion (about 30% of chrIII) spans the region from *MAT* to HMR, or, in one case, the region from *MAT* to *HML* (∼125 kb, or about 40% of chrIII). Clones with this deletion segregated 2 live:2 dead spores upon dissection of tetrads, consistent with the presence of multiple essential genes present on the deleted segment.

**Figure 2.**
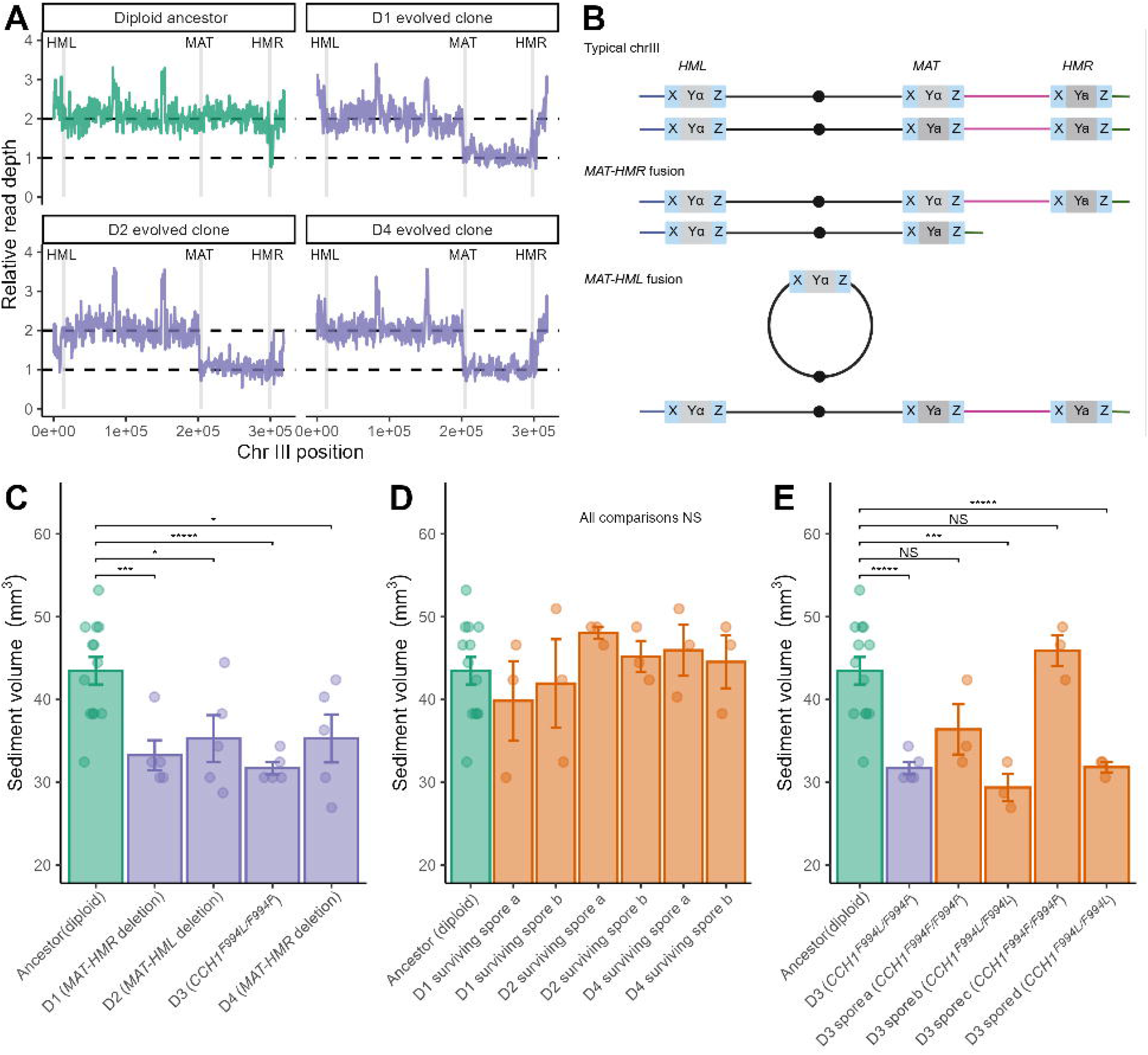
(A) Read depth in chromosome III showing *MAT-HMR* and *MAT-HML* deletions in clones isolated from evolved diploid populations. (B) Schematic of the products of *MAT-HMR* and *MAT-HML* fusions. (C) Clone isolated from diploid populations have reduced sedimentation compared to the ancestor. (D) Dissection products of evolved diploid clones cured of their deletion have the ancestral sedimentation phenotype. (E) Dissection products of the *CCH1* heterozygote show that the reduced-sedimentation phenotype segregates with the F994L homozygote.

Whole-population sequencing of populations D1, D2, and D4 at Day 57 revealed that the *MAT-HMR* or *MAT-HML* deletion was fixed or nearly fixed. We also found evidence for the presence of deletion lineages in 3 of 4 Day 29 populations D5, D6, and D8 (Figure S5 in File S1), so that in total a *MAT-HMR* or *MAT-HML* deletion of observable frequency occurred in 6 of 8 diploid populations evolved under selection for increased cell separation.

The *MAT-HMR* deletion is historically known as Hawthorne’s deletion, and the *MAT-HML* deletion is known as Strathern’s circle (Hawthorne 1963; Strathern et al. 1979; Herskowitz 1988). The latter deletion (or equivalently, fusion) is expected to produce a ring chromosome III (Figure 2B). The arrangement of one linear and one circular chrIII is not expected to be deleterious under mitotic growth, but is expected to depress spore viability because crossovers during meiosis between the ring and the linear homolog generate a dicentric chromosome (Haber et al. 1980; Haber et al. 1984). While we did not observe any reduction in spore viability below 50% (Table S1 in File S1), the following evidence points to the existence of a true ring chromosome III: (1) a PCR product spanning *MAT* and *HML* is produced; (2) we performed long-read sequencing and found approximately the expected number of long reads mapped to regions left of *MAT* and right of *HML* (Figure S3 in File S1); and (3) neither short-nor long-read sequencing suggested a new breakage site.

### The association between large deletions and an increased cell separation phenotype

Evolved clones with either the *MAT-HMR* or *MAT-HML* deletion exhibited significantly less sedimentation than the ancestor (Figure 2C), and an analysis of the distribution of cell cluster size, as quantified by microscopy, showed that these evolved strains were less clumpy than the ancestor (Figure 3A). For example, the ancestral proportion of cells occurring in clusters of 4 cells or fewer was 43%; for the Hawthorne’s deletion strains, this estimate ranged from 65% to 77%.

**Figure 3.**
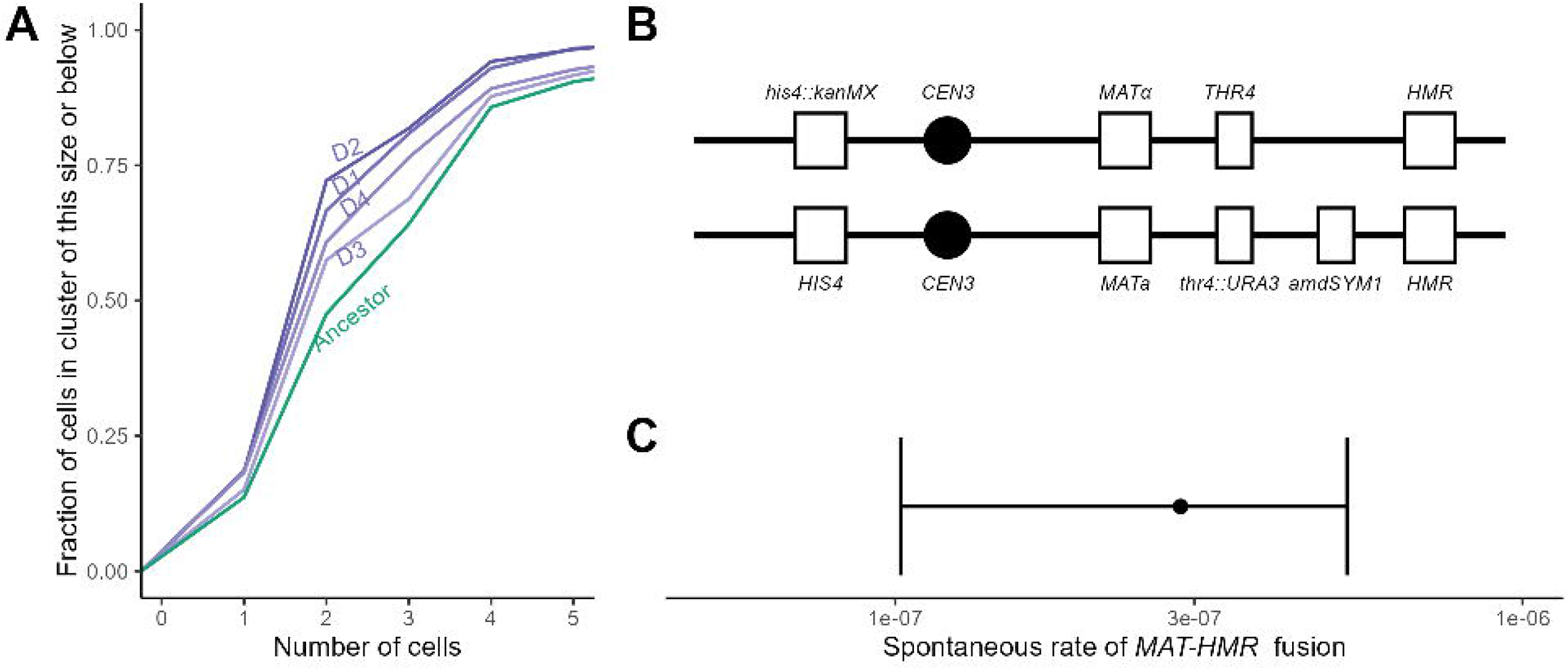
(A) The empirical cumulative distribution of cell cluster size for evolved diploid genotypes. (B) Selection scheme to enable estimation of the rate of Hawthorne’s deletion: one copy of chrIII was doubly tagged with counterselectable markers between *MAT* and HMR, while complete loss of chrIII was selected against by histidine auxotrophy. (C) The spontaneous rate of *MAT-HMR* deletion of one copy of chrIII as estimated by the fluctuation assay.

We also quantified the budding pattern of the evolved clones. Diploid yeast tend to bud from alternating poles (Chant and Pringle 1995). We observed growing cells microscopically and scored whether two consecutive buddings occurred on alternate poles, and found a significant depression in the frequency of alternate-pole budding for clones with the *MAT-HMR* or *MAT-HML* deletion isolated from the evolved populations, compared to the ancestor (Table S3 in File S1). This more disordered budding may be connected to the increase in cell separation.

Besides the parallelism observed in the evolution experiment, three additional lines of evidence support a causal relationship between the *MAT-HML/HMR* fusions and an increased cell separation phenotype. First, seeking to rule out that these large deletions were unconditionally beneficial in the experimental environment, we carried out a control evolution experiment under the same conditions as the main experiment, except that there was no selection against settling. In these populations, we observed no sign of Hawthorne’s deletion (Figure S5 in File S1). Second, we cured evolved clones of their deletions. The live products of dissected tetrads from the *MAT-HML* or *MAT-HMR* deletion clones, being homothallic, yielded diploid colonies cured of the deletion. These cured diploids no longer showed the reduced-sedimentation phenotype displayed in the strains with the deletion (Figure 2D). Third, as described below, we measured the rate of the *MAT-HMR* fusion in the diploid ancestor. Spontaneous clones isolated from this experiment, possessing the *MAT-HMR* fusion, produced significantly less sediment during settling than the ancestor (Figure S7 in File S1), akin to the evolved clones.

### The rate of Hawthorne’s deletion

The time to fixation of a mutation under positive selection depends on both the strength of selection and the rate of mutation. For rare mutations, there may be a substantial waiting time and even when the mutation occurs, selection does not act deterministically until the lineage is of a certain size. The observed parallelism in this experiment suggested that the per-generation rate of Hawthorne’s deletion could not be too small. To confirm this hypothesis we estimated the spontaneous rate of Hawthorne’s deletion via the classic fluctuation assay. We used genetic modifications (Figure 3C) to facilitate selection for *MAT-HMR* deletions while counter-selecting against the complete loss of chrIII. We found a spontaneous rate of loss of *MAT-HMR* of 2.9e-07 (95% CI: 4.1e-08 to 3.4e-07). This rate represents the generation of *MAT-HMR* fusions from a single marked copy of chrIII, so the total rate of *MAT-HMR* fusions is twice as large, and the total rate of any Hawthorne’s deletion including *MAT-HML* fusions is, by implication, up to four times the measured rate, although we did not measure the rate of *MAT-HML* fusion.

### A dominant point mutation to CCH1 also produces less clumpy growth in diploids

Diploid population D3 did not have a high-frequency Hawthorne’s deletion. The representative clone that we isolated had only one called mutation, a nonsynonymous heterozygous mutation (F994L) to the calcium channel gene *CCH1*, which is located on chrVII. Population sequencing revealed this mutation fixed or at high frequency (Figure S6 in File S1). The representative isolated clone sedimented significantly less than the ancestor (Figure 2C) and was quantitatively less clumpy (Figure 3A). We sporulated and dissected tetrads of this isolate to produce F994L/F994L and F994F/F994F homozygotes, the genotypes of which were confirmed by Sanger sequencing. The F994F/F994F homozygotes did not show a reduced sedimentation phenotype, while the F994L/F994L homozygotes displayed a sedimentation phenotype very similar to the Hawthorne’s deletion strains (Figure 3E). The sedimentation phenotype of the F994L/F994L homozygote is not more extreme than that of the F994L/F994F heterozygote, implying that this mutation is dominant with respect to cell separation. We noticed when carrying out dissections that F994L colonies were smaller than F994F colonies, suggesting that under this experiment’s particular selection regime, the heterozygote has the highest total fitness considering both growth rate and selection for cell separation.

*CCH1* is homologous to human *CACNA1A* (Paidhungat and Garrett 1997) and encodes the voltage-gated calcium channel subunit α1A. Variants of *CACNA1A* are implicated in several neurological diseases (Rajakulendran et al. 2012). Alignments of yeast *CCH1* and human *CACNA1A* show that yeast F994 is located on a transmembrane helix and homologous to human F708. A search of ClinVar (Landrum et al. 2014) revealed one observation (accession: RCV000622937.4) of human F708L, reported in association with developmental and epileptic encephalopathy, a severe epilepsy syndrome.

## Discussion

### Selection on AMN1 in haploids was predictable

Amn1 causes clumpy growth in haploids by mediating the degradation of Ace2, the major transcription factor of the post-mitotic cell separation program (Fang et al. 2018). It is likely that semi-inadvertent selection for *AMN1* mutants is part of the evolutionary history of common lab strains, since cultures of well-separated cells are more experimentally tractable for various purposes. This experiment has in effect recapitulated that selection; thus it is not surprising that we recovered many *AMN1* mutants, although we did not observe the particular *AMN1* allele, 368V, that is present in S288C and related strains. It is a demonstration of the usefulness of experimental evolution in genotype-to-phenotype mapping, if a selection gradient relevant to the phenotype of interest can be devised, that had *AMN1*’s role in governing cell separation in haploids not been previously known this experiment would have clearly pointed towards it.

### The Rim101 pathway is a novel regulator of cell separation during vegetative growth

Mutations to the Rim101 pathway were less expected. This pathway is known to have a role in pH sensing (Yan et al. 2020) and regulation of cell size (Shimasawa *et al*. 2023), and to connect to pathways regulating filamentous growth (Cullen and Sprague 2012). Arras *et al*. (2022) found that Rim101 pathway genes are required for sexual aggregation, but a role in regulation of cell separation during typical vegetative growth conditions has not previously been reported. We suspect that the clear effect of Rim101 pathway genes on cell separation reported here has previously escaped notice because many laboratory strains such as S288C have a nonfunctional allele of *AMN1* and are therefore already well-separated, which masks the increased cell separation resulting from disabling a Rim101 pathway gene in a background with functional *AMN1*.

The cell wall of yeasts requires specialized structures to effectuate cell division. At the mother-bud neck, septins provide a diffusion barrier and scaffold to help organize septum-building enzymes. The primary septum, made of chitin, acts as a temporary wall between daughter and mother cell that is subsequently degraded from the daughter cell by the chitinase *Cts1* (Weiss 2012). Ace2, which as noted above is degraded via Amn1, activates transcription of *CTS1*. Interestingly, *Δrim101* and *Δrim13* mutants have been shown to have increased transcription of *CTS1*, although there does not appear to be direct repression of *CTS1* by Rim101, and *Δrim101* mutants have an asymmetric distribution of the septin Cdc3 and an abnormal localization pattern of Chs4, a recruiter of chitin synthase III (Lamb and Mitchell 2003). These data suggest two possible non-exclusive mechanisms through which loss-of-function Rim101 pathway mutants might cause increased cell separation: increased production of chitinase, and production of an altered septum more vulnerable to degradation. Rim101 pathway mutants and *AMN1* mutants do not have identical phenotype in our assays: Rim101 pathway mutants produce more sediment (Fig 1C), but have a smaller settling distance (Fig 1D), compared to *AMN1* mutants. The latter finding is consistent with the notion that Rim101 pathway knockouts have decreased cell size (Shimasawa et al. 2023), given that smaller cells are expected to settle more slowly.

Interestingly, whole-population sequencing revealed that at least some populations maintained competing *AMN1* and Rim101 pathway lineages even after 57 days of evolution (Table 1). It is possible that certain uncontrolled aspects of our experimental design—e.g. the precise depth of the pipet tip used to transfer part of the culture after the settling period—influenced whether *AMN1* or Rim101 pathway mutants predominated in any particular haploid population. Strain and mating type differences could also have played a role, although we do not have the statistical power to assign any such effects.

### Implications of the rate of large deletions involving the mating type loci

In diploids, we observed repeated selection for *MAT-HMR* or *MAT-HML* deletions on chromosome III. These heterozygous large deletions (roughly 30-40% of chrIII) arise as a consequence of homology shared by the three mating type loci. They are historically known as Hawthorne’s deletion and Strathern’s circle, respectively, and were instrumental in the working out of the mating type switching system (Hawthorne 1963; Strathern et al. 1979; Haber et al. 1980; Herskowitz 1988; Haber 1998). Hawthorne’s deletions were initially observed as a small proportion of rare mating-type switching events in heterothallic (*ho*) strains. In this respect, it is interesting that we have observed these deletions in homothallic (*HO*) wild-type diploids under vegetative growth, that is, separately from any obvious context of mating type *switching* per se.

Our point estimate of the rate of *MAT-HMR* fusion on a single marked copy of chrIII, 3.5e-07 per generation, implies, if the rate of *MAT-HML* fusion is roughly equivalent, a total rate of either deletion on the order of 1e-06 per generation. This estimation was performed in ho diploids, by necessity of the process of strain construction; although *HO* is silenced in *MAT*a/*MAT*α diploids it is possible that leaky expression of *HO* could drive a higher rate if some *HO*-initiated events resolve as deletions. The estimated rate is one to two orders of magnitude higher than the mutation rate to loss of function for a typical gene (Lang and Murray 2008). As such, it is conceivable that the observed parallelism of *MAT*-HMR/*HML* deletion is the result of selection for haploinsufficiency of a single gene, but we think it is more likely that gene dosage effects at least two loci in the span of the deletion are involved.

As these large deletions may be considered segmental aneuploidies or monosomies, it is also illustrative to compare to rates of whole-chromosome aneuploidy. In that context, the measured rate is less than the average per-chromosome rate of aneuploidy observed in Zhu *et al*. (2014) and Sharp *et al*. (2018) (∼1e-04/16 = ∼6e-06), but higher than the average per-chromosome rate of reduction in copy number reported in those studies, the large majority of observed aneuploidies (∼90%) in both being gains in copy number.

In nature, *MAT-HMR/HML* deletions are presumably selected against as they are lethal in the haploid state. Therefore the rate of such deletions may be considered to be a kind of load imposed by the three-cassette mating-type system. As this rate is roughly comparable to rates of aneuploidy, this load appears tolerable (although as discussed below if aneuploidy often provides transient adaptive states, then aneuploidy rates are not true loads). Assuming that the rate of *MAT-HMR/HML* fusion is indeed a load in natural populations, the long-term benefits of the mating-type system must necessarily outweigh this and other loads associated with sexual reproduction.

### Aneuploidy as adaptive mechanism

Several previous evolution experiments have observed aneuploidy in *S. cerevisiae* and other yeasts in response to various selective pressures (Gresham et al. 2008; Rancati et al. 2008; Selmecki et al. 2009; Chen et al. 2012; Gilchrist and Stelkens 2019). Most often, an increase in chromosomal copy number is observed, but there are some reports of selection for monosomy (Yang et al. 2013; Barney et al. 2021). It is hypothesized that adaptation via aneuploidy might be important in natural populations by providing a transient adaptive state, especially as natural isolates of *S. cerevisiae* tolerate aneuploidy more easily than lab strains (Muenzner et al. 2024 May 22). It would be interesting to extend the evolution reported here to observe how and if this segmental aneuploidy is resolved into, or replaced by, some euploid adaptive genotype, or whether it persists indefinitely.

### Conclusions and future directions

In this experiment, we selected for increased cell separation in haploid and diploid populations of wild yeast strains for over 380 generations. In haploids we found parallel selection for loss-of-function mutations to *AMN1* and for loss-of-function mutations to genes of the Rim101 pathway, pointing to a previously unrecognized role for the Rim101 pathway in regulating cell separation. In diploids, we found parallel selection for large heterozygous deletions spanning *MAT-HMR* or *MAT-HML* on chromosome III. These large deletions caused less sedimentation, smaller clusters of cells, and an altered budding pattern. We also found one diploid population fixed for a heterozygous mutation to *CCH1*, a calcium channel known to play a role in emergence from mating pheromone-induced arrest (Fischer et al. 1997) and other stress responses that depend on calcium influx. Further work is required to elucidate the roles of *CCH1* and the Rim101 pathway in influencing cell separation, and to determine how heterozygous *MAT-HMR/HML* deletions affect this phenotype. Continued evolution in this system could lead to insights regarding the long-term maintenance of large heterozygous deletions in diploids and/or the role of transient states of aneuploidy in adaptation.

## Supporting information

All supplemental figures and tables

All mutations observed

## Data availability statement

File S1 contains all supplementary figures and tables. File S2 is a spreadsheet of all called mutations observed in the evolved clones. Raw sequencing reads for this study, including both ancestors, and evolved clones and populations, have been deposited in the NCBI Sequence Read Archive (SRA) under BioProject accession <u>PRJNA1397233</u>. Ancestral and evolved strains are available upon request.

## Acknowledgements

We would like to thank Erfei Bi and Hiroki Okada for kindly sharing relevant deletion strains from the yeast deletion collection.

