## Supplementary material for "Directed evolution for cell separation in natural isolates of budding yeast reveals selection to deactivate *AMN1* and the Rim101 pathway in haploids and selection in favor of Hawthorne’s deletion in diploids": All supplemental figures and tables

#### Supplemental Figures

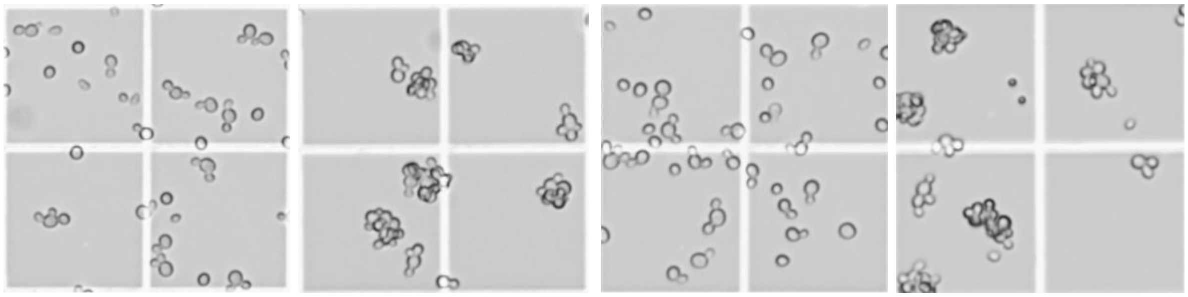

$AMN1^{Cys452Tyr}$ $\Delta AMN1^{Cys452Tyr}::AMN1^{Anc}$ $AMN1^{Leu472Pro}$ $\Delta AMN1^{Leu472Pro}::AMN1^{Anc}$

**Figure S1.** In evolved clones, when the derived *AMN1* allele is replaced with the ancestral allele, the clumpy phenotype is restored.

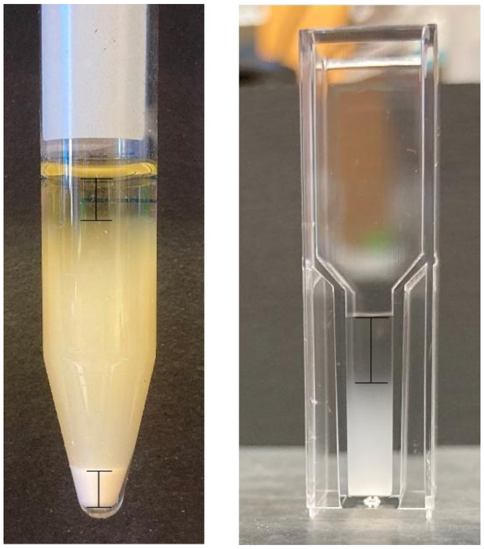

**Figure S2.** Sample photographs of sedimentation and settling distance assays as described in Methods. Photographs not to same scale.

A

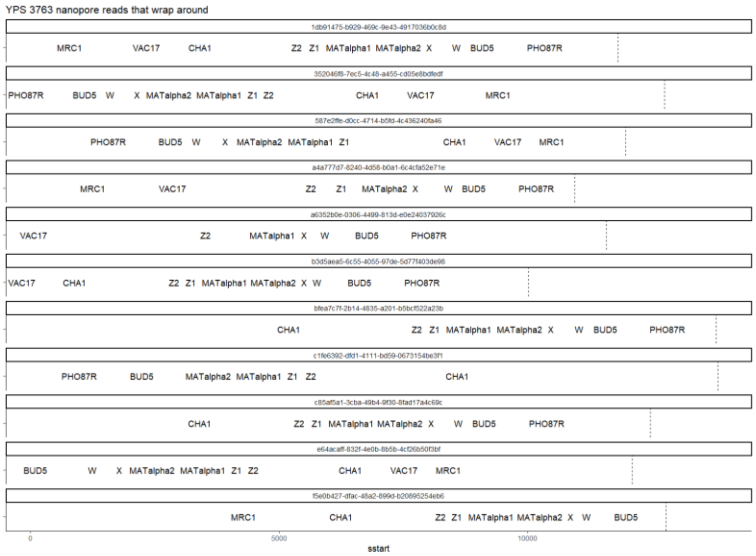


B

| Reads >= 10 kb | |
|---|---|
| Spanning HML | 7 |
| Spanning MAT | 18 |
| HML-MAT fusion | 11 |

**Figure S3.** Long-read (Nanopore) sequencing evidence of a *MAT-HML* fusion on chr III from the clone isolated from population D2. (A) Individual long reads with BLAST hits to genes right of *HML* (*MRC1*, *VAC17*, *CHA1*) and left of *MAT* (*BUD5*, *PHO87*) indicated. (B) The number of long reads that support an *MAT-HML* fusion is about equal to the number of long reads that canonically span *HML* or *MAT*, consistent with the clone being heterozygous for the Hawthorne's deletion.

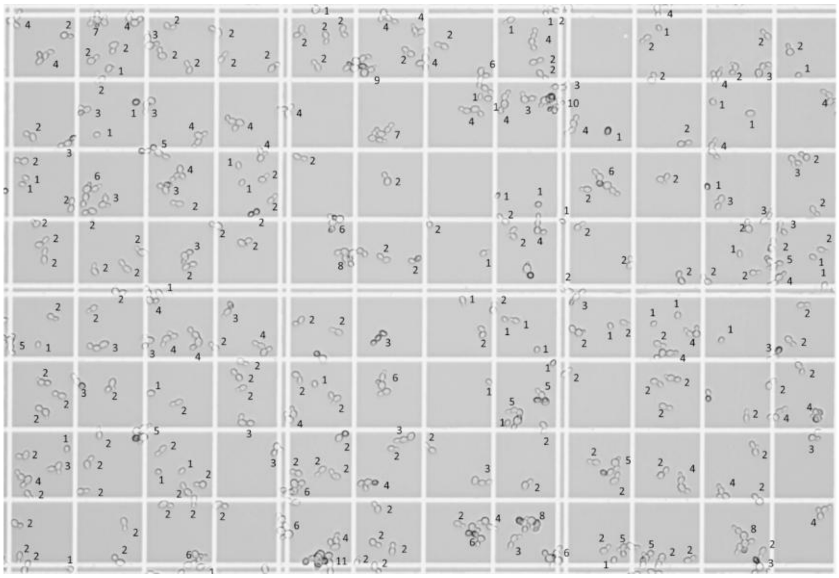

**Figure S4.** A sample image of a cultured of a diploid clone after labelling of cluster size. Data generated in this manner underpins the empirical CDF shown in Fig. 3a.

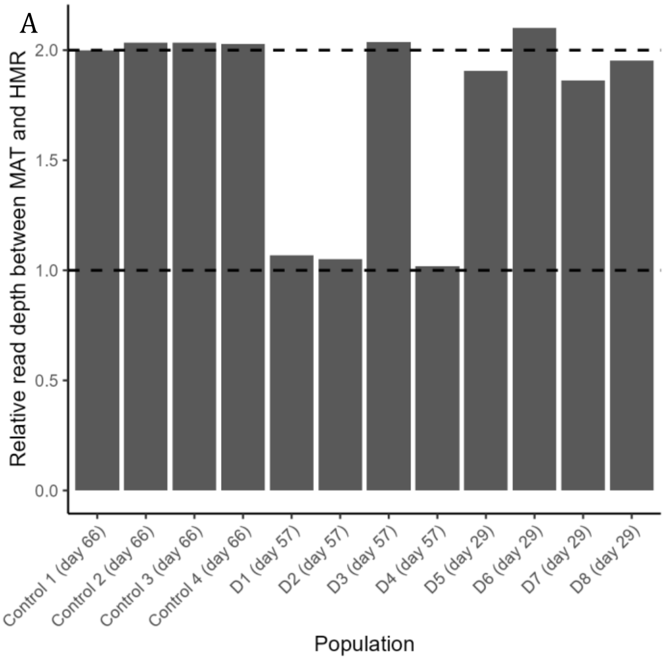


B

| Population | Clones checked | Tetrads segregate 2:2 live:dead | Tetrads normal |
|---|---|---|---|
| D5 (day 29) | 10 | 6 | 4 |
| D6 (day 29) | 10 | 0 | 10 |
| D7 (day 29) | 10 | 2 | 8 |
| D8 (day 29) | 10 | 5 | 5 |

**Figure S5.** (A) The median read depth of the *MAT-HMR* region of chrIII, normalized by the median read depth of the *HML-MAT* region and scaled so that the ancestral value is 2, for 12 evolved diploid populations. Three of the four diploid Day 57 populations (D1, D2, D4) are fixed or nearly fixed for Hawthorne's deletion (or Strathern's circle); three of the four diploid populations halted at Day 29 (D5, D7, and D8) show evidence of a low-frequency Hawthorne's deletion; and the control (no selection against clumpy growth) populations show no evidence of  a Hawthorne's deletion. (B) Dissection results: 5, or sometimes 10, tetrads from each of 10 clones isolated from each of the D5-D8 diploid Day 29 populations were dissected and scored for normal spore viability or 2:2 live:dead. The dissection results agree qualitatively with the read depth data that there are Hawthorne's deletion lineages in D5, D7, and D8.

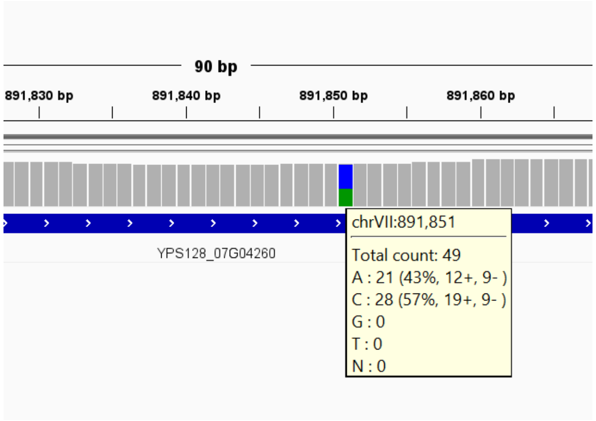


**Figure S6.** Screenshot from IGV showing the heterozygous *CCH1* mutation Phe994Leu fixed or nearly fixed in population D3 at Day 57.

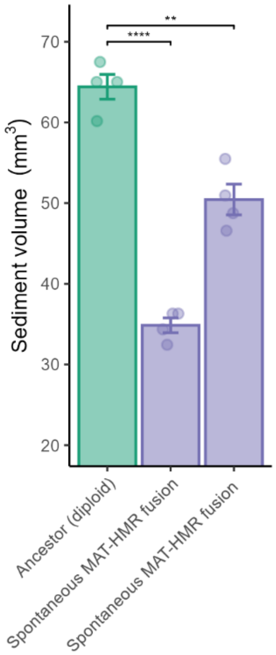


**Figure S7** Two spontaneous *MAT-HMR* fusion clones isolated from the fluctuation test show significantly less sedimentation than the ancestor. Culture growth for this assay was carried out in SD + uracil, as these clones are uracil auxotrophs.

### Supplemental Tables

| | Surviving spores per tetrad | | | | |
|---|---|---|---|---|---|
| | **1** | **2** | **3** | **4** | **Total viability** |
| D1 clone (*MAT*-*HMR* fusion) | 1 | 9 | 0 | 0 | 0.48 |
| D4 clone (*MAT*-*HMR* fusion) | 0 | 13 | 0 | 8 | 0.69 |
| D2 clone (*MAT*-*HML* fusion) | 1 | 18 | 2 | 0 | 0.51 |

**Table S1.** Spore viability of the diploid chrIII deletion isolates from the evolution experiment.

| Replicate | Colonies on plate | Colonies picked | Of picked: sporulated | Of sporulated: segregated 2:2 live:dead | Of 2:2 segregators: positive PCR for MAT-HMR fusion | Adjusted count |
|---|---|---|---|---|---|---|
| 1 | 413 | 32 | 29 | 0 | NA | NA |
| 2 | 38 | 38 | 2 | 2 | 1 | 1 |
| 3 | 57 | 32 | 17 | 6 | 6 | 11 |
| 4 | 32 | 32 | 7 | 7 | 7 | 7 |

**Table S2.** Raw data for the fluctuation test measuring the rate of spontaneous MAT-HMR deletion. Replicate #1 is a “jackpot” for a non-Hawthorne’s deletion and it was therefore not possible to count the true number of colonies with a Hawthorne’s deletion. For replicate #3, the count of Hawthorne’s deletion colonies was adjusted accounting for the fraction of colonies examined, i.e. (57/32) * 6 = 11 (rounded to the closest integer).

| | Alternating pole | Other pattern | *p*-value differs from ancestor |
|---|---|---|---|
| Diploid ancestor | 65 | 31 | -- |
| D1 clone (*MAT*-*HMR* fusion) | 55 | 51 | 0.03 |
| D2 clone (*MAT*-*HML* fusion) | 51 | 57 | 0.005 |
| D3 clone (CCH1 F994L/F994F) | 52 | 33 | 0.4 |
| D4 clone (*MAT*-*HMR* fusion) | 50 | 55 | 0.004 |

**Table S3.** The evolved clones with a chrIII deletion have a significantly different budding pattern from the ancestor, while the evolved clone with a *CCH1* mutation does not. Fisher’s exact test was used for calculation of *p*-values.
